# Differentiation of white matter histopathology using b-tensor encoding and machine learning

**DOI:** 10.1101/2023.02.17.529024

**Authors:** Ricardo Rios-Carrillo, Alonso Ramírez-Manzanares, Hiram Luna-Munguía, Mirelta Regalado, Luis Concha

## Abstract

Diffusion-Weighted Magnetic Resonance Imaging (DW-MRI) is a non-invasive technique that is sensitive to microstructural geometry in neural tissue and is useful for the detection of neuropathology in research and clinical settings. Tensor valued diffusion encoding schemes (b-tensor) have been developed to enrich the microstructural data that can be obtained through DW-MRI. These advanced methods have proven to be more specific to microstructural properties than conventional DW-MRI acquisitions. Additionally, machine learning methods are particularly useful for the study of multidimensional data sets. In this work, we have tested the reach of b-tensor encoding data analyses with machine learning in different histopathological scenarios. We achieved this in three steps: 1) We induced different forms of white matter damage in rodent optic nerves. 2) We obtained *ex-vivo* DW-MRI with b-tensor encoding schemes and calculated quantitative metrics using Q-space Trajectory Imaging. 3) We used a machine learning model to identify the main contributing features and built a voxel-wise probabilistic classification map of histological damage. Our results show that this model is sensitive to characteristics of microstructural damage. In conclusion, b-tensor encoded DW-MRI analyzed with machine learning methods, have the potential to be further developed for the detection of histopathology and neurodegeneration.

## Introduction

Non-invasive inference of tissue microstructure is made possible through diffusion-weighted magnetic resonance imaging (DW-MRI) [1]. This technique has been useful to characterize cerebral connectivity, plasticity, development, and diverse pathologies. The need to find clinical standardized DW-MRI biomarkers in healthy and pathological neural tissue has driven more research in this field [2, 3]. Classical DW-MRI techniques (i.e., those encoding diffusion through a single pair of pulsed gradients) have shown sensitivity to nervous tissue damage but not specificity to diverse histopathological forms [3]. Multidimensional diffusion encoding (MDE) DW-MRI [4] techniques were developed to address this situation. Specifically, the b-tensor encoding technique [5] provides a robust framework to explore multidimensional diffusion data. The theoretical background of these techniques is robust, and they have been tested in controlled environments with simulations [6] or in healthy tissue [7].

One of the main advantages of using MDE DW-MRI acquisitions is that the complex information in the data is adequate for advanced diffusion models or signal representations. In the diffusion tensor distribution (DTD) [4] a collection of micro-diffusion tensors with different shapes, sizes, and orientations describe microstructure. Such complex micro-structural models are not attainable through standard DW-MRI acquisitions. Thus, MDE DW-MRI can potentially characterize certain neuropathological changes in detail. However, relatively few studies have used this technique for this purpose [8].

## Materials and methods

### Animals

We used adult male Wistar rats for this study (weight: 354 ± 59 g). Animals were held in a vivarium room under normal light/darkness conditions with controlled temperature and humidity. Animals had *ad libitum* access to food and water. The study was approved by the Bioethics Committee of the Institute of Neurobiology, Universidad Nacional Autonoma de Mexico (protocol 096.A) under NOM-062-ZOO-1999 law. All procedures were performed in compliance with ARRIVE guidelines.

### Animal surgery

Normal rats were used to investigate two forms of white matter pathology: axonal degeneration and inflammation (Fig 1). Rats were anesthetized with a ketamine/xylazine mixture (70mg/kg and 10mg/kg ip) and placed on a well-illuminated surface. For each animal, the procedure was as follows: the right optic nerve was lesioned while the left one remained intact. This allows a direct comparison between subjects and between groups. Rats were divided into four different groups:

1. Axonal degeneration (n=6). Induced through unilateral retinal ischemia [9]. Animals were placed in a stereotaxic frame. A 32-gauge needle was inserted into the anterior chamber of the right eye of each rat, and connected to a reservoir with saline solution that was elevated until an in-line pressure monitor indicated 120 mmHg (higher than systolic pressure); this pressure was maintained for 90 min.
2. Inflammation (n=9). Elicited through injection of *1μl* of lipopolysaccharide (LPS, 4.5 *μg/μl;* Sigma-Aldrich) in the optic nerve [10]. A small lateral incision behind the eye was performed. Then, lacrimal glands and extra-ocular muscles were dissected to expose the optic nerve. Using a 32-gauge needle coupled to a Hamilton syringe, the injection was done approximately 1 cm rostral to the optic chiasm. After careful and slow manual injection, the needle was left in place for approximately 1 minute in order to avoid reflux. The skin was sutured and topical antibiotics were administered. Animals were allowed to recover from anesthesia and placed in their cages until perfusion.
3. Saline solution injection (n=9). This group aimed to evaluate the mechanical damage produced by the sole needle insertion. The procedure was identical as the previous group but the injection consisted of 1 *μl* of saline solution.
4. Control (n=8). Healthy animals with both optic nerves intact.

**Fig 1.**
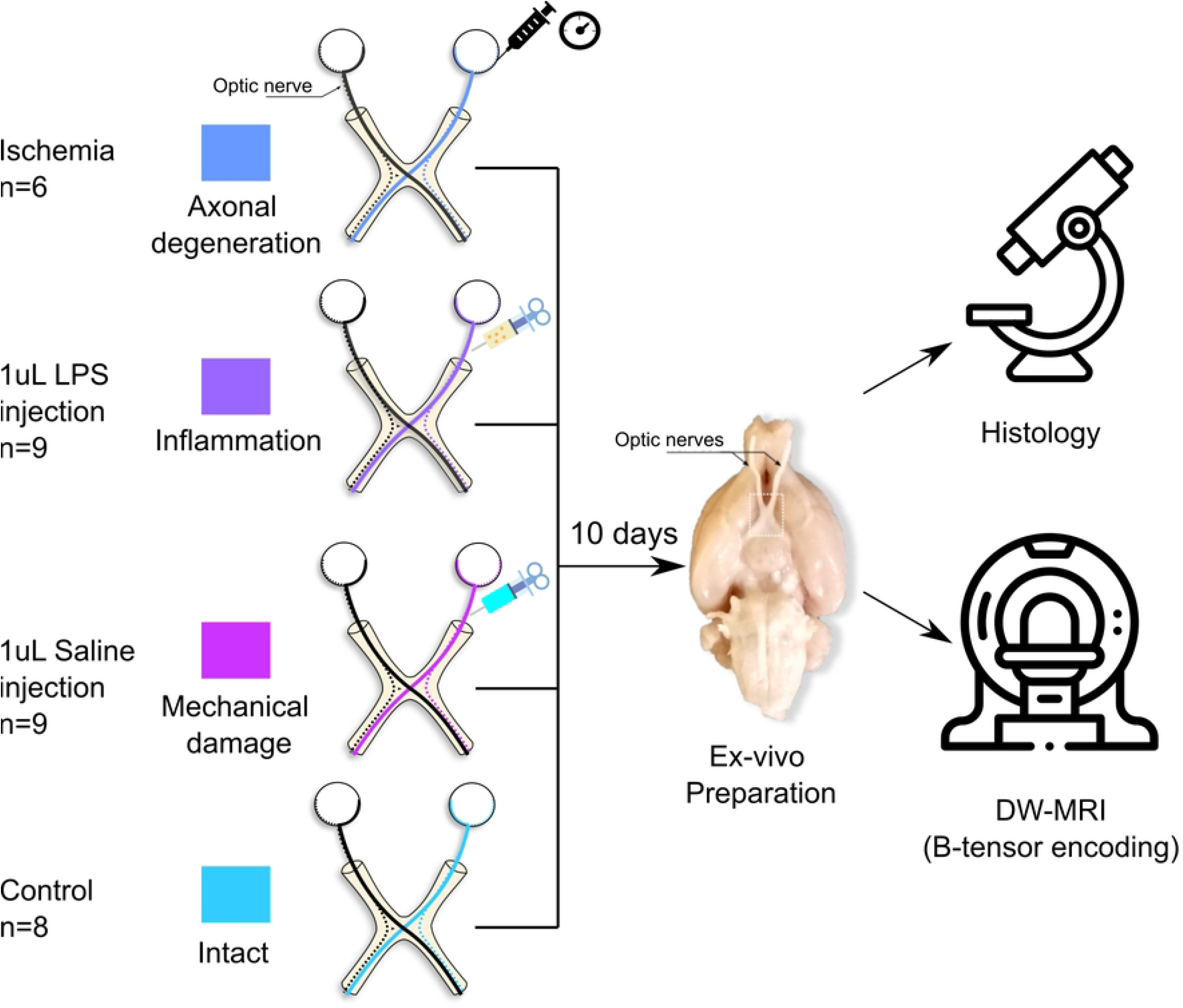
Experimental design. Axonal degeneration or inflammation of the right optic nerves was induced *in vivo* through retinal ischemia or LPS injection, respectively. Additionally, saline solution was injected into a group of animals to evaluate mechanical damage. Animals were sacrificed ten days after experimental procedures, tissue was fixed, and the brains and optic nerves were extracted. b-tensor encoding DW-MRI were acquired *ex vivo*.

### Brain extraction

Ten days after the surgical procedure, all the animals were deeply anesthetized using an intraperitoneal overdose of sodium pentobarbital. Animals were transcardially perfused with 0.9% sodium chloride followed by paraformaldehyde(4%)-glutaraldehyde (2.5%) solution. Brains were carefully extracted leaving at least 1 cm of optic nerves intact. Specimens were post-fixed in fresh 4% paraformaldehyde solution at 4 °C until scanning day.

### Imaging

Brains were scanned 15 ± 10 days post-extraction. The most distal portions of the optic nerves were attached to the ventral side of the olfactory bulbs by using cyanoacrylate in order to prevent the optic nerves from floating during the scan. To achieve a reduced field of view for DW-MRI, we carefully dissected and kept the basal portion of the brain. These specimens were immersed in Fluorinert (FC-40, Sigma-Aldrich) and allowed to rest for four hours at room temperature before scanning. Acquisition protocols were carried out at the National Laboratory for Magnetic Resonance Imaging using a 7 T Bruker Pharmascan with 760 mT/m gradients and a Cryoprobe. The scanning room temperature was 21 ± 1 °C, and the Cryoprobe’s heated ceramic head mount was set at the same temperature. DW-MRI images were acquired using the available sequence in the Preclinical Neuro MRI repository (https://osf.io/ngu4a); which is based on a 2D spin-echo sequence. Voxel resolution was 80 × 80 × 1000 μm^3^. Other MRI parameters: TR = 1500 ms, TE = 30.9 ms, two averages, flip angle = 79°, scan time = 16 h.

DW-MRI were obtained with b-tensor encoding based on a previously described protocol [7]; specific modifications were done for our *ex vivo* setting. The protocol consists of 3 different gradient waveforms to obtain linear, planar, and spherical tensor encodings (LTE, PTE, and STE, respectively). STE and PTE waveforms were optimized and Maxwell-compensated [11] using NOW toolbox [12]. LTE waveforms were extracted from the optimized STE waveforms to obtain similar gradient spectral characteristics between waveforms [13]. All waveforms have the same duration (*δ*_1_ = 9.8, δ_2_ = 10.4, separation time = 5.72 ms), and each one was scaled in gradient magnitude to achieve 4 different b-values (0.5, 1.4, 2.8 and 4 ms/μm^2^). The STE waveform was rotated to obtain 10 directions for every b-value. Rotating the STE waveforms results in the same spherical b-tensor, but this redundancy ensures a more robust data processing [7]. LTE and PTE waveforms were rotated to obtain [10,10,16,46] directions for each corresponding b-value. S1 Fig shows the waveforms and protocol used in this experiment.

### Image data preprocessing

Given the long spin-echo based acquisition, the obtained images do not present many artifacts. The only preprocessing step needed was denoising, as the high b-value shells (4 ms/μm^2^) are noisy, which we achieved through Marčenko-Pastur PCA [14,15] as implemented in *mrtrix3* [16]. Examples of final images for each encoding acquisition are shown in S2 Fig. Regions of interest (ROI) for injured and control optic nerves were manually drawn in 3 to 4 slices per nerve (92 ± 25 voxels for each nerve).

### Analysis of b-tensor encoded DW-MRI

We fit the QTI method to the obtained b-tensor encoding images and extract eight microstructural metrics. Four of them capture the macroscopic behavior of the DTD ensemble and are akin to those from diffusion tensor imaging (DTI) [17]: 1. Fractional Anisotropy (FA). 2. Mean diffusivity (MD). 3. Axial diffusivity (AD). 4. Radial diffusivity (RD).

The following four QTI metrics capture the microscopic behavior of the DTD ensemble and are only achievable through methods such as b-tensor encoding:

5. Micro Fractional Anisotropy (μFA). Measures the mean value of all the fractional anisotropy values of all tensors in the DTD.
6. Orientation coherence (*C_c_*). Measures the level of orientation coherence of the micro tensors in the DTD.
7. Isotropic kurtosis (*K_i_*). Quantifies the kurtosis produced by the size variance of the micro tensors in the DTD.
8. Anisotropic kurtosis (*K_a_*). Quantifies the kurtosis produced by the microscopic anisotropy.

We obtained QTI metrics using the implementation in QTI+ [18]. The original QTI implementation is biased to very complex microstructure [6], while QTI+ provides a more stable solution to the DTD fitting optimization problem and achieves smoother and more precise maps than the standard QTI implementation. We used the default settings for QTI+. To avoid regions where DTD fitting was poor, we excluded voxels (6.8% of all data) where any of the QTI metrics resulted in values outside their valid range: Normalized metrics (FA, *μ*FA) should lie between 0 and 1, and kurtosis metrics (*K_i_* and *K_a_*) should be between 0 and 5.

### Histology

Following dMRI acquisition, specimens were returned to 4% paraformaldehyde solution and kept at 4°C until processing. Briefly, the optic nerves were separated from the basal portion of the brain and were washed with buffered sodium cacodylate (0.1 M) and glutaraldehyde (3%). Then, stained with osmium tetroxide (0.1%), washed with cacodylate buffer (0.1 M), and dehydrated with ethyl alcohol at different concentrations (10%, 20%, 30%, until absolute). Next, samples were embedded in a 1:1 epoxy resin/propylene oxide solution for 42 h. For polymerization, samples were placed in a plastic container with epoxy resin and kept at 60°C for 36 h. Finally, each block was sectioned (600 nm thick) using an ultramicrotome (RMC PowerTome PT XL). Slices were stained with a toluidine blue/sodium tetraborate solution (both 5%).

### Histology images

Photomicrographs were obtained with a Leica DM750 microscope (equipped with a 5M pixels digital camera) with x10 and x100 objectives, and an Amscope T690C-PL microscope (equipped with a 10M pixels digital camera) with a x40 objective. We transformed the images to 16-bit grayscale and digitally enhanced their contrast using Fiji [19] (version = 2.9.0). Images with the x40 lens were stitched using the stitching plug-in [20] available in Fiji.

### Machine learning pipeline

Visual inspection of the photomicrographs revealed histological patterns that overlapped between experimental groups (see Histological evaluation). We therefore chose to (i) reframe the classification labels for the machine learning pipeline into histological classes that reflect different types of histopathological damage, as Intact, Injured, and Injured+, and (ii) analyze voxels of the regionally-affected nerves, identified as the Regional pattern (see Fig 2).

**Fig 2.**
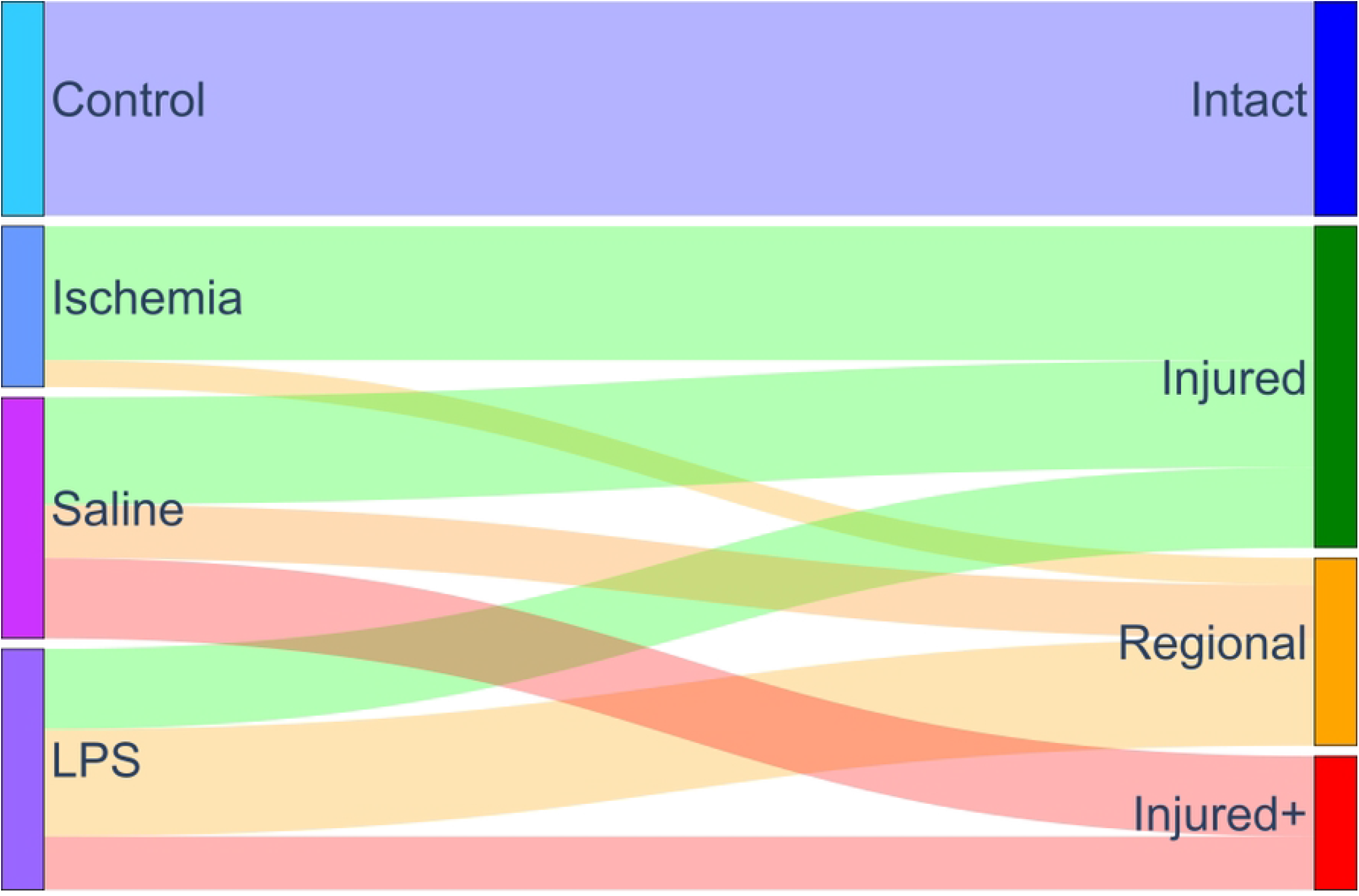
Labeling system based on histological patterns. Classes were assigned to each nerve after a visual examination of histology, based on the spatial pattern and type of histological characteristics. The left column represents the experimental procedures, while the right column indicates the labels used for the identification of tissue type based on diffusion properties.

Fig 3 shows a diagram with the machine learning (ML) pipeline. QTI+ data from Intact, Injured, and Injured+ classes were used for the train/test set in the ML pipeline in a voxel-wise fashion (A). We trained a random forest model [21] (B) (80/20% fold) and conducted a feature relevance analysis by Gini importance [21] with scikit-learn (version=1.1.2, https://scikit-learn.org). We classified each voxel in the Regional nerves with this model (C and D). The resulting probability of class membership is visualized as a composite red-green-blue (RGB) map (E), with each channel representing each tissue class: Intact:Blue, Injured:Green, and Injured+:Red.

**Fig 3.**
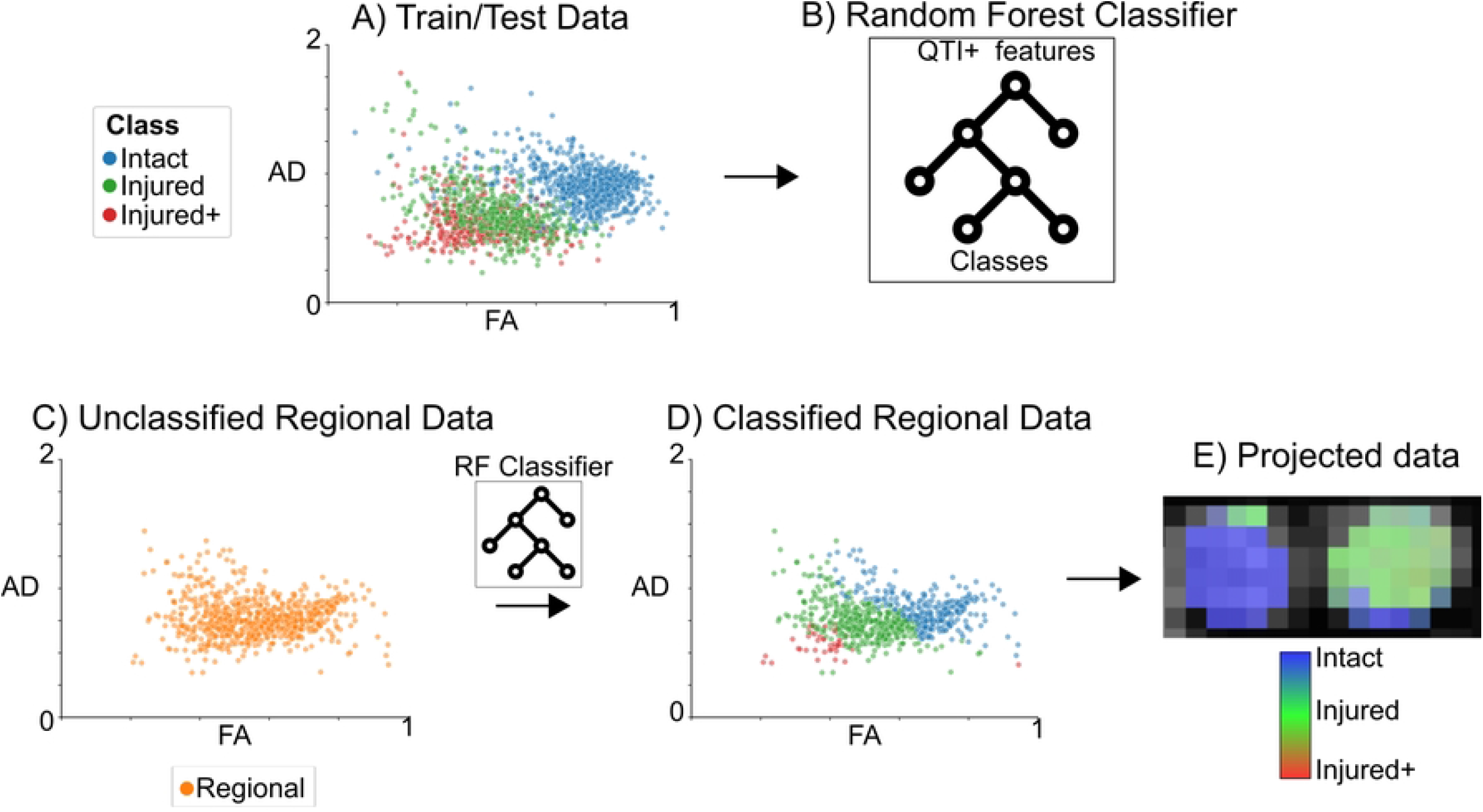
Diagram of the machine learning pipeline. We used the QTI+ data for all voxels labeled according to histology (Panel A: each color-coded data point represents a voxel) as input to train the random forest model (B). We classified each voxel of the regionally-affected nerves (Regional) (C) into histological damage classes (D). Finally, we projected the classified data back into an anatomic RGB map that quantifies tissue damage (E).

Feature relevance analysis is a complex subject with potential caveats. Previous work indicates that Gini importance has two main problems: First, it tends to be biased towards features with high cardinality [22]. This, however does not apply to our data since it is on a continuum. Second, Gini importance reports statistics related to the training set [21]. Thus, we also performed a feature relevance analysis by permutations on the test set [21]. After we report the accuracy/F1-Score results and feature analysis with the test set, we calculated a bootstrapped estimator to estimate the variance of the permutation feature analysis and checked if they maintained the same order of relevance. To this end, we randomly permuted the train/test partitions to perform 200 different experiments (using the same optimized hyperparameters reported for the random forest model) to evaluate the reproducibility of the permutation feature relevance analysis. We emphasize that this analysis is done after the main analysis with the train/test set that is reported in the Results (Machine learning classification), and its only purpose is to check for biases of feature analysis related to the original train/test partition.

### Data availability

All DW-MRI data is available through the Open Science Framework (https://osf.io/b2k4z/).

## Results

### Experimental labels for DW-MRI data

Quantitative maps derived from QTI+ showed great asymmetry between the intact and affected nerves for most of the metrics (Fig 4) caused by the lesion. This was confirmed by analysis of the average values per nerve. Fig 5-A,B shows the per-animal average difference between the intact (left) and experimental nerve (right), indicating large differences of FA, AD, μFA and *C_c_* between the two nerves. Diffusion metrics from nerves in the three experimental conditions showed considerable overlap between them but were clearly different from the intact nerves (Fig 5-C, D). S3 Fig shows the overall distribution by experimental groups.

**Fig 4.**
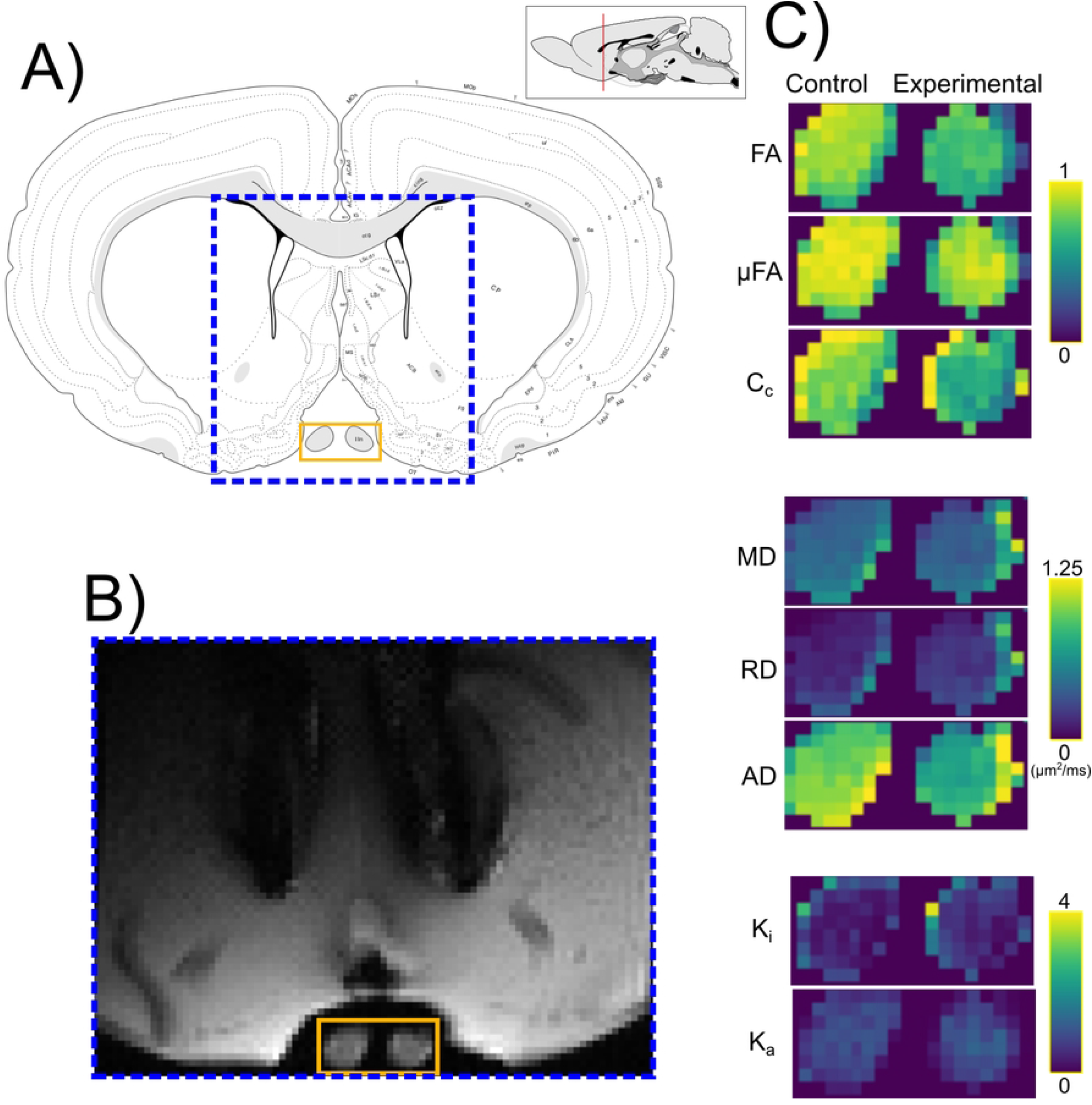
Q-space Trayectory Imaging contrasts. A) Anatomical atlas reference (adapted from [23]). DW-MRI were obtained from the portion of the brain specimen indicated by the dashed blue box. B) Example denoised DW-MRI with spherical b-tensor encoding (b=2.8 *ms/μm*^2^). C) Enlarged images corresponding to the orange rectangle in panel B. QTI metrics for control (left) and experimental (right) optic nerves. (Abbreviations: fractional Anisotropy (FA), microscopic fractional anisotropy (*μ*FA), orientation coherence (*C_c_*), mean diffusivity (MD), radial diffusivity (RD), axial diffusivity (AD), isotropic kurtosis (*K_i_*) and anisotropic kurtosis (*K_a_*)).

**Fig 5.**
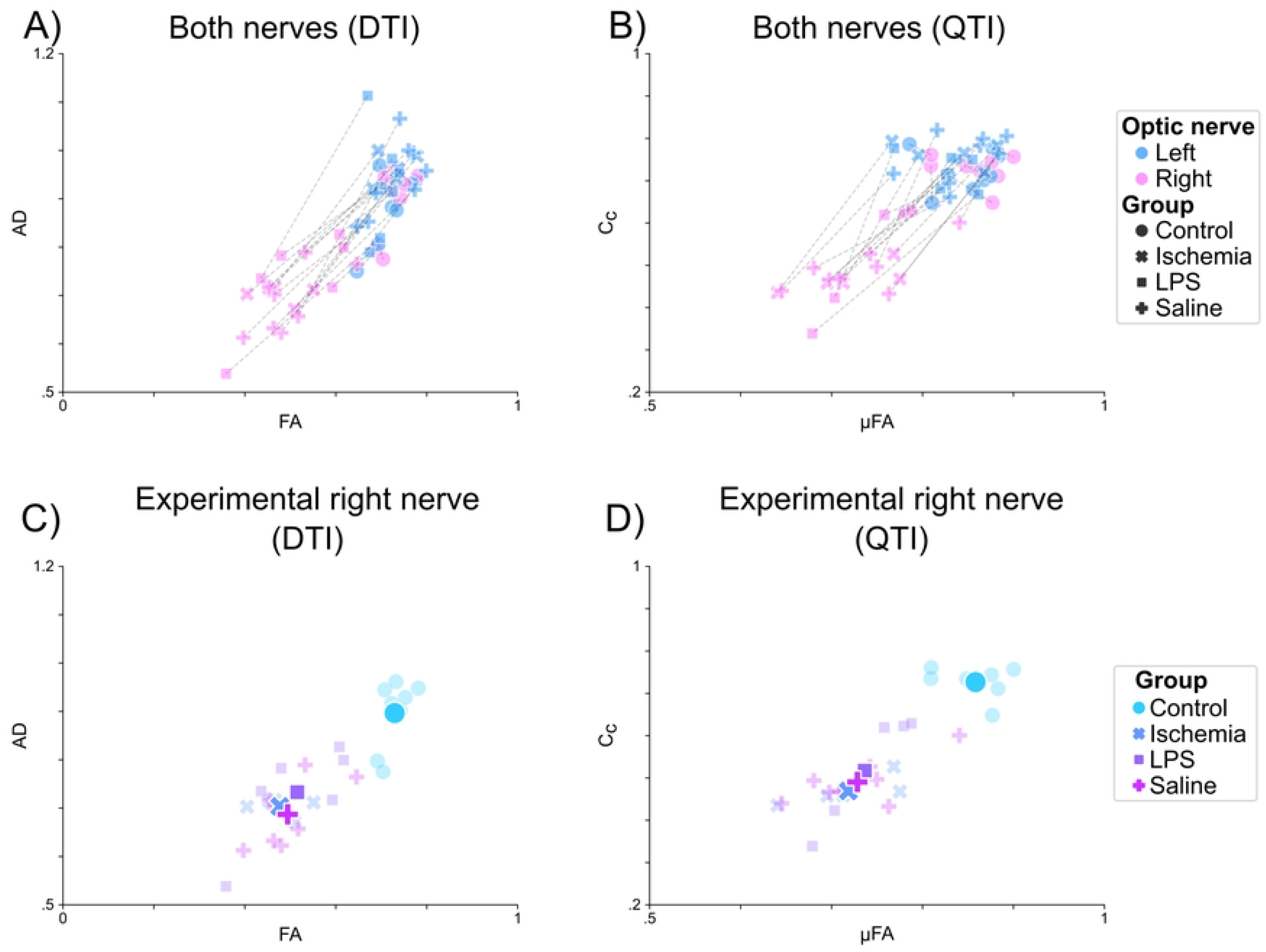
QTI metrics by experimental group. Data points correspond to the average values of all voxels of each optic nerve, per subject. A,B: Intact (Left) vs Experimental (Right) optic nerves. Lines connect the two optic nerves of each animal. C,D: Right optic nerves color-coded according to experimental procedure. Semi-transparent markers show average values per animal; average values for each experimental condition are indicated as large solid markers.

### Histological evaluation

Histological examination of sections stained with toluidine blue (see Histology) showed that retinal ischemia induced diffuse axonal degeneration and mild gliosis. Nerves injected with LPS also had reductions of axonal density and more glial cell infiltration. There was evidence of independent mechanical damage, as injections of saline solution showed a range of axonal degeneration and gliosis ranging from mild to severe. Moreover, while retinal ischemia induced tissue injury mostly in a spatially-homogeneous fashion, nerves treated with either type of injection produced damage either homogeneously or only within confined regions of the nerve, with some areas showing damage, and others displaying nearly intact structure. Thus, we manually labeled each nerve based on the type and spatial pattern of histopathology, as (1) *Intact*, (2) *Injured*: characterized by globally-reduced axonal density; (3) *Injured+:* displaying homogeneous and profound axonal loss and severe gliosis; and a (4) *Regional* pattern, with different regions of the nerves showing either of three histological types (Fig 2). This classification system allowed us to perform a spatial assessment of microstructural damage produced by the experiments, i.e., we used the diffusion properties of the Intact, Injured, and Injured+ classes to identify the corresponding histological patterns in the regionally-affected nerves. Photomicrographs in Fig 6 show examples of the histopathological patterns identified. Panel A is a prototypical *Intact* nerve, characterized by a large number of axons with clearly-defined myelin sheaths and bright intra-axonal space, interspersed with angular glial cell processes. Panel B shows an *Injured* nerve, displaying reduced axonal density, numerous collapsed axons with dark intra-axonal space (green arrow), and reactive glial cells with large, amoeboid processes. Panel C shows an *Injured+* nerve, nearly devoid of axons with a considerable amount of glia in a reactive foamy state. Lastly, panel D shows a Regional nerve, with large and clearly-delimited regions that can be described with the three aforementioned classes. As noted in (Fig 2, this classification based on the histological type was used to perform voxel-wise ML-based classification from QTI+ metrics.

**Fig 6.**
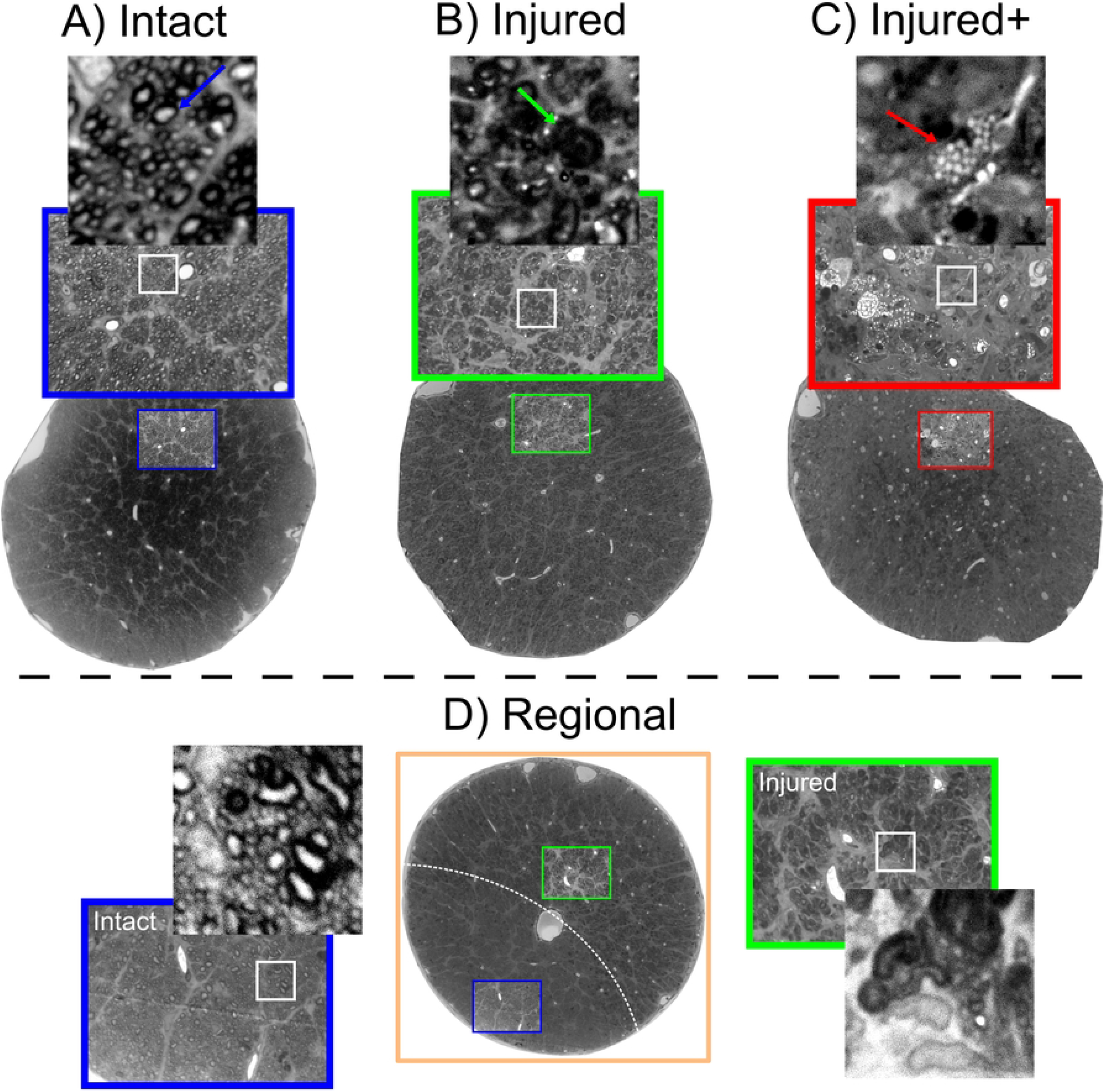
Histopathological patterns after experimental procedures. A: Intact nerve with a large number of axons and narrow glial processes. B: Injured nerve with collapsed axons (green arrow), reduced number of axons, and gliosis. C: Injured+ nerve with very few axons and large reactive glial processes with foamy interior indicative of myelin degradation (red arrow). D: Regional nerve showing clearly separated areas (dashed white line) of either of the three histological patterns. The areas in the regional nerves have characteristics of the intact and injured classes, making them a suitable fit for a machine learning classification problem. Photomicrographs of whole nerves acquired at x10 magnification; photomicrographs in colored squares acquired at x100 magnification.

### Histology-based labels for DW-MRI data

A voxel-wise inspection according to the histology-based classes of the right (experimental) nerves reveals differences in the distributions for QTI metrics in groups (Fig 7). Albeit their large overlap, it is possible to visually separate the distributions. Group-wise analyses showed considerable alterations of QTI metrics in all histopathological types, characterized by reduced FA, AD, *μ*FA, and *C_c_*, and increased RD and *K_i_* (S4 Fig). MD showed slight reductions in the Injured and Injured+ conditions.

**Fig 7.**
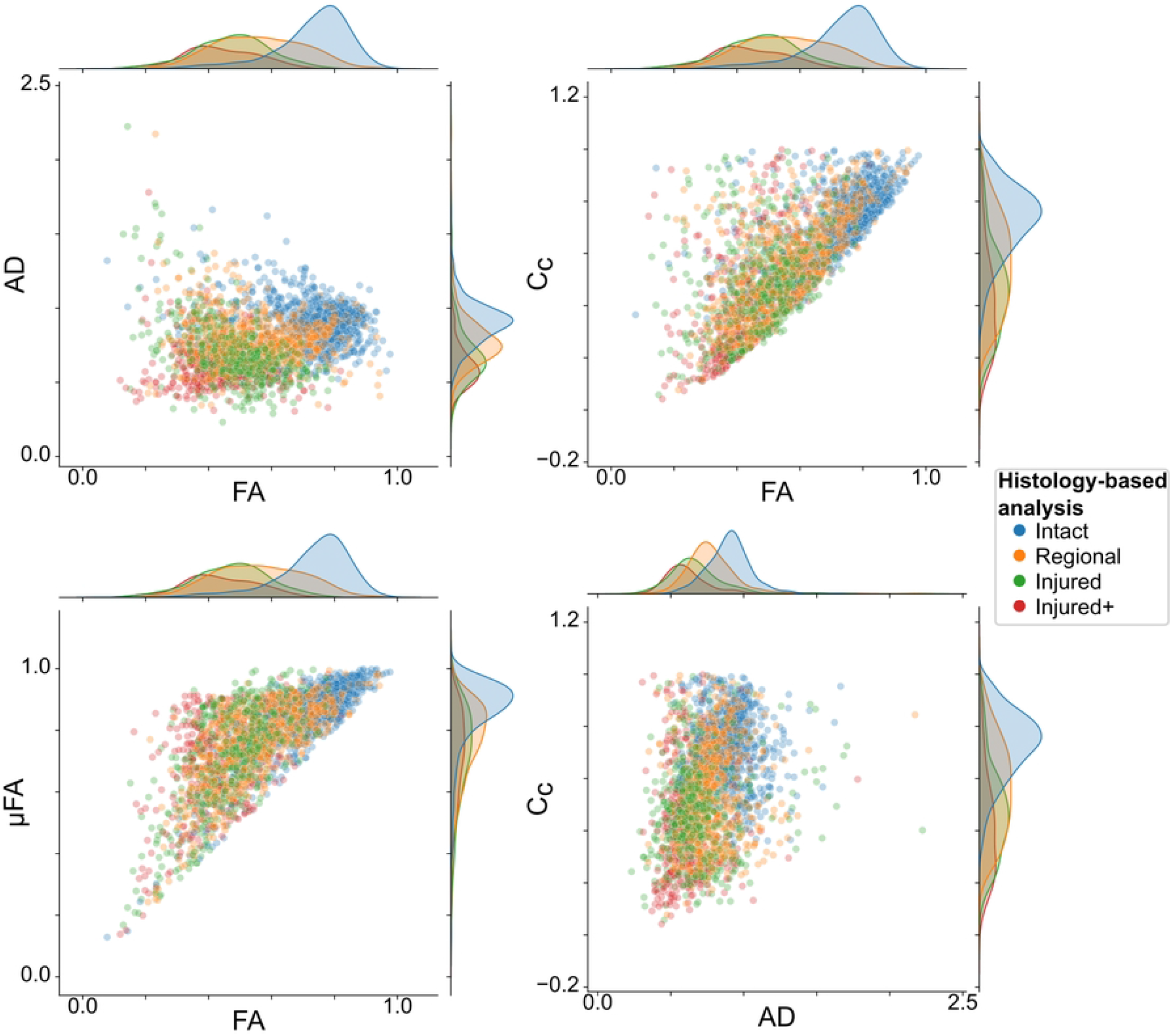
Voxel-wise scatter plots according to the histology-based labeling system. Metrics from the Intact class (blue) are clearly different from those of the experimental classes. Metrics from Injured and Injured+ classes are overlapped but still separable. The Regional class, being composed of areas of either Intact or any of the two injured classes, shows diffusion metrics distributed across the metrics space.

### Machine learning classification

We trained a random forest model for the voxel-wise classification of histopathological classes in nerves identified as having Regional abnormalities, according to the pipeline in Fig 3 (maximum depth=6; number of estimators [trees]=100). The overall classification accuracy was 80.11% and an F1-score of 79.4% (with a weighted average for multiclass classification) to distinguish between the three histopathological classes. Fig 8A shows the confusion matrix for the classification of the test data set. Fig 8B shows the results of the feature relevance analysis. FA and AD—metrics from the classic DTI—are the two most relevant features for the machine learning model. S5 Fig shows the permutation feature analysis and the bootstrapped feature analyses, which confirmed the relevance of FA, AD and *C_c_* for classification, in that order.

In addition to illustrating the classification pipeline, Fig 3D shows voxels from Regional nerves (Fig 3C) classified with the ML method. Fig 3E shows an example of classified voxels as a RGB map. The majority of voxels within left nerves (intact) are correctly classified (blue–intact). The right (experimental) nerves show most of the voxels identified as Injured, with spatial patterns that correspond to histology (see Fig 6).

**Fig 8.**
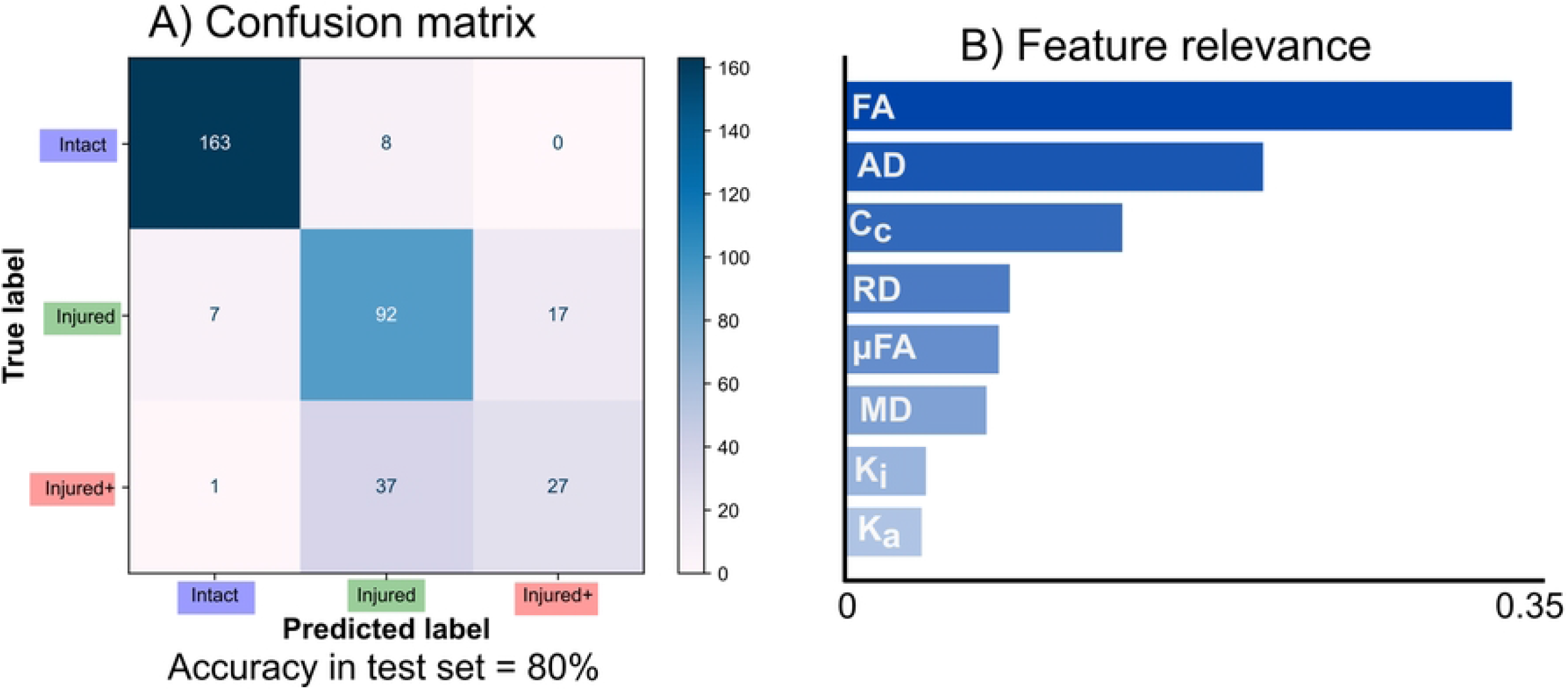
Machine learning results. A: Confusion matrix for classification in the test set. B: Feature relevance from the random forest. FA and AD, both derived from DTI, are highly important for classification. With the exception of *C_c_*, metrics derived from QTI are less relevant.

An example voxel-wise classification of a single nerve (LPS injection and identified as Regional, see Fig 2) shows the algorithm is sensitive to microstructural degeneration. (Fig 9). Voxels identified as Injured and Injured+ are larger in number in rostral slices (i.e., nearest to the injection site), with more caudal slices gradually showing more voxels classified as Intact. Notably, photomicrographs of the same nerve at the approximate same levels as the DW-MRI show a similar spatial pattern of injury and corresponding histopathological pattern as that identified by the random forest. The vast majority of voxels in the left (intact) nerves are correctly identified as Intact. S6 Fig shows three more examples of correct histopathological damage classification.

**Fig 9.**
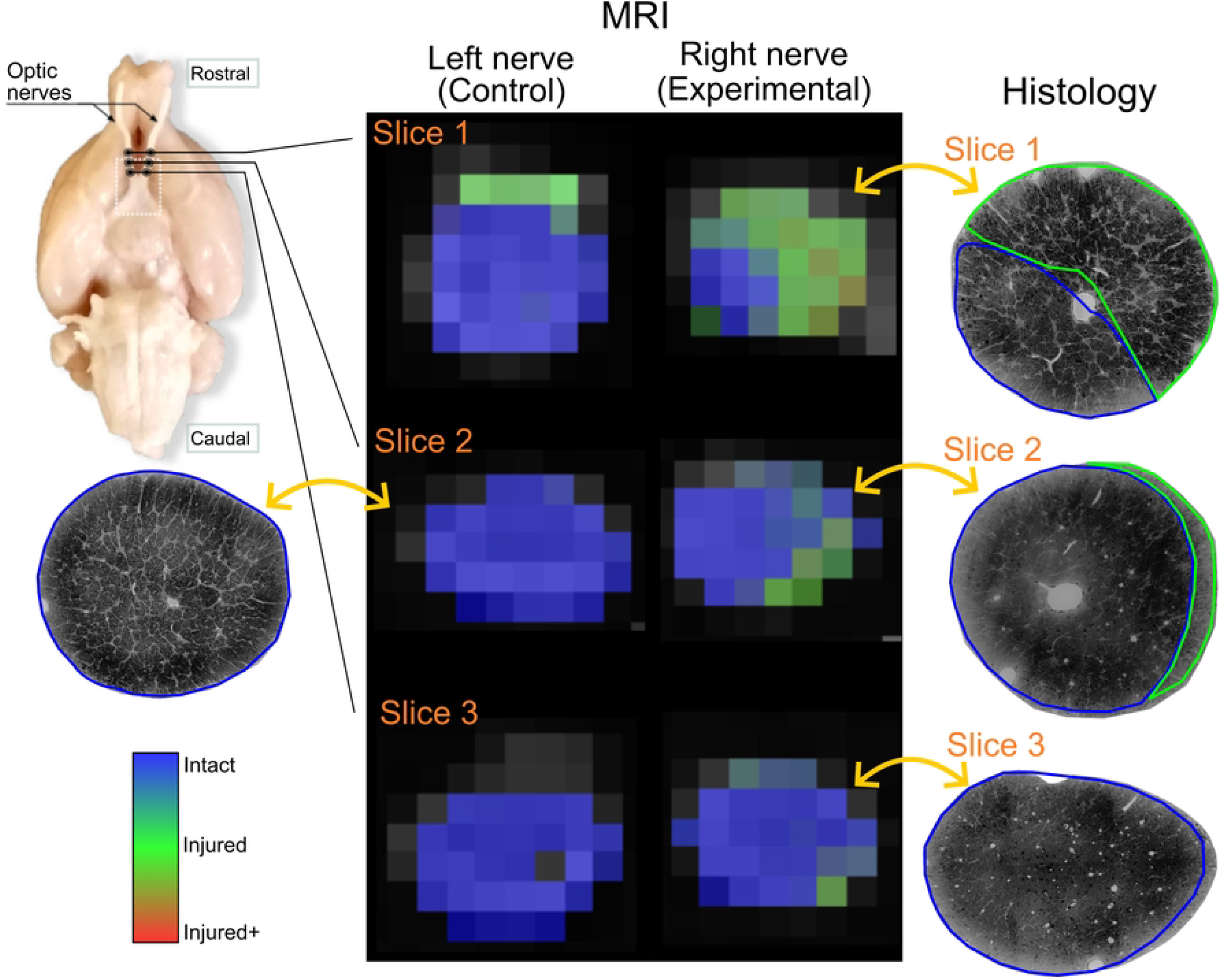
Voxel-wise classification of histological patterns. Rat histological example data showing Regional damage of the left and right (experimental) nerves. The two optic nerves are shown in three different slices in rostro-caudal order. Photomicrographs of the same experimental nerve at the approximate same locations show clearly demarcated areas of Injured and Intact histological patterns, that correspond to the voxel-wise classification of the DW-MRI of the experimental nerve.

## Discussion

In this work, we explore the synergy of QTI metrics and machine learning for the non-invasive identification of white matter histological damage. Our data show that these metrics are sensitive to altered histological patterns. Two DTI and one QTI metrics were the most relevant for accurate classification of tissue damage. i.e. the metric achievable only through b-tensor encoding further improved the results obtained from the machine learning pipeline.

The optic nerve has been widely used to evaluate white matter changes through DW-MRI. A common approach is to induce retinal ischemia that results in Wallerian degeneration of the retinal ganglion cells and their axons throughout the lesioned nerve, which reflects as specific patterns of diffusion abnormalities [24,25]. In this study, we explored the possibility to detect and differentiate between inflammation and axonal degeneration. Considering that both processes present with different severity in several neurological disorders, the aim is to achieve better methods to differentiate them and improve the diagnostic yield of dMRI. However, no tissue damage differentiation was observed between optic nerves only mechanically damaged and those injected with LPS; both showed axonal degeneration and gliosis at diverse severity levels. Injections of the nerve produced enough tissue damage to reduce FA, similar to the reductions caused by retinal ischemia (Fig 5) [24,25]. Decreased μFA was also observed in all experimental conditions, indicative of an increase of isotropic diffusion profiles from glial cells present as a result of inflammation and mechanical tissue damage, as confirmed by histology (Fig 9). Dispersion, as seen with Cc, was also reduced in all affected nerves, which fits the observed tissue disorganization of the experimental nerves. The rest of the metrics derived from DTI or QTI overlapped among all the experimental conditions (Fig 5C,5D). This prevented us from establishing a clear differentiation between inflammation and axonal degeneration.

We observed that mechanical damage varied from subtle axonal degeneration to the annihilation of the entire axonal population (Fig 6). We therefore, re-classified our data based on histological findings and their spatial extent (Fig 2), with the injured nerves (Injured and Injured+) distinguished by the presence of foamy reactive glia that usually appears only in advanced stages of degeneration [26, 27]. In addition, many injected nerves showed a mosaic of Intact, Injured, and Injured+ histopathologies, which we set out to automatically classify based on the diffusion profiles derived from nerves with spatially homogeneous tissue characteristics. As we were working with complex 8-dimensional data from thousands of voxels, this was an ideal setting for a machine learning application.

Random forest models were used based on the following facts: 1) Less prone to overfit, 2) easier to interpret (it’s possible to inspect and interpret the individual estimators-i.e. decision trees-in the model), 3) the variance in the estimators provides resilience to: (i) noise and (ii) poor quality data points, 4) feature relevance analysis is straightforward. We obtained similar results when using state of the art machine learning methods like XGBoost [28] and neural networks [29]. This indicates that classification performance is more related to the nature of our data than to the classification algorithm used.

The overall accuracy performance of the automatic classification was high (80%). While the best distinction performance was between Intact and the two Injured classes, there was a modest success in the differentiation between the Injured and Injured+ classes (Fig 8A). Confusion between the two degrees of injury may be due to axonal loss (present in both types) acting as the main microstructural characteristic driving the measured diffusion properties. Other dMRI modalities specific to glial cells [30] or combined with other MRI modalities like spectroscopy [31] could disentangle these cases.

Feature relevance analysis (Fig 8B) revealed that FA and AD are the most relevant features to differentiate between tissue types. This was expected, as both are sensitive to the overall loss of anisotropy in white matter capturing the main effect of degeneration. DTI metrics are sensitive but not specific. We expected features exclusive to B-Tensor encoding to improve the classification algorithm by providing additional information, given their specificity to certain properties of microstructure [8]. For instance, we expected *C_c_* to capture the increased axonal dispersion typical of WM degeneration, as previous studies using standard DW-MRI acquisitions had suggested [32]. Indeed, *C_c_* is the third important feature in the analysis. Nevertheless, Cc is (by QTI definition) correlated to FA, and therefore contributes less to the classification problem if FA is already included in the analysis. Indeed, repeating the same pipeline using only QTI features revealed that *C_c_* is the most relevant feature of the analysis while preserving a similar classification performance (not shown). μFA was only slightly relevant for the classification; we hypothesize that gliosis reduces μFA in a similar pattern in all experiments, thus reducing its efficacy as a predictor. While DTI metrics capture the overall loss of diffusion anisotropy, QTI metrics could be capturing the fine details in the diffusion properties to separate these conditions. We also expected *K_i_* (related to the variance of sizes in the DTD model [5]) to be increased as a result of glial infiltration. *K_a_* might explain the loss of micro anisotropy in the medium and is also related to axonal loss. The relatively low explained variance in the data by the kurtosis metrics may be attributed to the bias secondary to the assumption of the DTD model that *μ*K is equal to zero [33], which is not the case in degeneration [34], and therefore K_a_ and *K_i_* may both be absorbing this effect. Note that features achievable with DTI (FA and AD) capture the main properties of neurodegeneration in aligned white matter bundles, but dispersion (*C_c_*) and *μ*FA could be important factors in white matter regions with crossing fibers or gray matter.

There are some limitations in this study. First, the experimental procedures (particularly those related to direct nerve injections) produced overlapping histopathologies. This precluded the distinction between axonal degeneration and inflammation, and limits the interpretability of our findings. In particular, we cannot conclude from our data whether QTI is capable of resolving between those two histopathological processes. However, careful examination of histological slides allowed us to differentiate between Injured and Injured+ classes based on the presence of foamy glial cells and the extent of axonal loss, which were identified by the random forest algorithm based on diffusion metrics. Future work should try to minimize confounding factors introduced by mechanical damage of the tissue by utilizing other experimental approaches. Second, slice thickness was large (1 mm). Thick slices were acquired to improve the signal to noise ratio, but partial volume effects could introduce inaccuracies in the estimation of diffusion metrics, particularly for the Regional pattern as injured regions vary along the nerve. Third, STE and LTE waveforms were tuned [7,13] but this does not ensure they have the same diffusion time window [35]. Diffusion time dependence could be an important factor in neurodegeneration [3] and was not directly investigated or controlled for in this study; further studies should give some insight into the contribution of time-dependent diffusion to distinguish between types of histological damage. Last, machine learning applications benefit from large data sets. While our voxel-wise data set is not small the overall accuracy of the method could be improved with more data points.

There are other possibilities for the analysis of b-tensor encoding data. Like the diffusion tensor, QTI is a signal representation [36]. There are other interesting avenues of analysis like Diffusion Tensor Distribution imaging [37] that can extract direct DTD features or even extend it to multidimensional MRI analysis to capture relaxometry effects [38,39]. Approaches with biophysical models using b-tensor encoding [40,41] can be used to extract microstructural properties that cannot be obtained without strong modeling assumptions using single diffusion encoding acquisitions. Nevertheless, they are based on the standard model of white matter that is applicable for healthy tissue, and it is unknown whether it would be adequate for the detection of severe deviations (i.e., tissue damage) without modifications to the underlying assumptions. More work is needed to test if these approaches to DW-MRI could identify tissue damage with high sensitivity and specificity.

Machine learning methods provide a new paradigm to understand and use the advanced methods available in the DW-MRI field. Direct visualization of tissue type probabilities as a color map (Fig 9), while proof of concept, provides a straightforward qualitative assessment of the type of damage at every voxel. The combination of spatial specificity and the availability of quantitative diffusion metrics can be a powerful tool to evaluate and diagnose microstructural changes in neurological disorders.

## Conclusion

In this work, we explore the ability b-tensor encoding methods to detect and differentiate between distinct forms of white matter pathologies. Specifically, we explored the metrics derived from QTI using state of the art machine learning methods. The majority of QTI metrics are sensitive to microstructural changes induced by neuropathology. While classic DTI metrics were the most important features for the training phase in the machine learning algorithm, features exclusive to b-tensor encoding improved its precision.

## Supporting information

**S1 Fig. Protocol scheme.** Full protocol scheme and example waveforms (b=2.8*ms/μm*^2^) used in this study.

**S2 Fig. b-tensor encoding example images.** Example preprocessed DW-MRI acquired b-tensor encoding (b=2.8 *ms/μm*^2^) of a single slice from one representative animal in the retinal ischemia group. Linear, planar and spherical tensor encodings (LTE, PTE, STE) and a non-diffusion-weighted image (b=0 *ms/μm*^2^) are shown. The yellow rectangle indicates the optic nerves.

**S3 Fig. Violin plots for the experimental groups for each QTI metric.**

**S4 Fig. Violin plots for the three nerve classes and regional pattern (defined by histological examination) for each QTI metric.**

**S5 Fig. Feature relevance analysis** A) Permutation feature relevance analysis in the test set. B) Bootstraped permutation feature analysis. FA and AD are the most important features. Gini importance (B) showed *C_c_* as the third-ranking relevant feature.

**S6 Fig. Examples of Regional histological damage and corresponding machine learning classification based on MDE DW-MRI.**

## Acknowledgments

This work was funded by CONACYT (FC 1782) and UNAM-DGAPA (IG200117, IA200621). Ricardo Rios-Carrillo is a doctoral student from Programa de Doctorado en Ciencias Biomédicas, Universidad Nacional Autónoma de México (UNAM) and received fellowship 707266 from CONACYT. Imaging was performed at the National Laboratory for Magnetic Resonance Imaging (Conacyt, UNAM, CIMAT). We thank Dr. Juan Ortiz for his technical assistance related to the MRI scanner. Ma. Lourdes Palma-Tirado and staff at the Microscopy Unit of the Institute of Neurobiology provided valuable help during histological processing. We thank Dr. Jorge Larriva-Sahd for helpful discussions and Gema Martínez for guidance related to histological methods.

## Notes

### Competing Interest Statement

The authors have declared no competing interest.

